# Genetic variation associated with PPO-inhibiting herbicide tolerance in sorghum

**DOI:** 10.1101/2020.05.04.076620

**Authors:** Pragya Adhikari, Emma Goodrich, Samuel B. Fernandes, Alexander E. Lipka, Patrick Tranel, Patrick Brown, Tiffany M. Jamann

## Abstract

Herbicide application is crucial for weed management in most crop production systems, but for sorghum herbicide options are limited. Sorghum is sensitive to residual protoporphyrinogen oxidase (PPO)- inhibiting herbicides, such as fomesafen, and a long re-entry period is required before sorghum can be planted after its application. Improving sorghum for tolerance to such residual herbicides would allow for increased sorghum production and the expansion of herbicide options for growers. To investigate the underlying mechanism of tolerance to residual fomesafen, a genome-wide association mapping study was conducted using the sorghum biomass panel (SBP) and field-collected data, and a greenhouse assay was developed to confirm the field phenotypes. A total of 26 significant SNPs (FDR<0.05), spanning a 215.3 kb region, were detected on chromosome 3. The ten most significant SNPs included two in genic regions (Sobic.003G136800, and Sobic.003G136900) and eight SNPs in the intergenic region encompassing the genes Sobic.003G136700, Sobic.003G136800, Sobic.003G137000, Sobic.003G136900, and Sobic.003G137100. The gene Sobic.003G137100 (*PPXI*), which encodes the PPO1 enzyme, one of the targets of PPO-inhibiting herbicides, was located 12kb downstream of the significant SNP S03_13152838. We found that *PPXI* is highly conserved in sorghum and expression does not significantly differ between tolerant and sensitive sorghum lines. Our results suggest that *PPXI* most likely does not underlie the observed herbicide tolerance. Instead, the mechanism underlying herbicide tolerance in the SBP is likely metabolism-based resistance, possibly regulated by the action of multiple genes. Further research is necessary to confirm candidate genes and their functions.

## 1 Introduction

Weed infestation is a major crop production constraint. Herbicide application is a critical control strategy in most crop production systems, and modern agriculture relies heavily on herbicides for weed suppression. Unfortunately, a limited number of herbicides are available for sorghum, and the herbicide options for grass control are even lower [1]. Moreover, sorghum is sensitive to many commonly used herbicides and is sometimes injured by herbicides labeled for use in sorghum [2]. Wet and poor soil physical conditions, delayed crop emergence, deep planting, seedling diseases, and poor-quality seed favor herbicide injury [2]. Thus, weed management in sorghum is challenging.

In recent years, protoporphyrinogen oxidase (PPO)-inhibiting herbicides have increased in popularity for the weed management of field crops. PPO-inhibitors were commercialized more than 50 years ago, but the introduction of transgenic glyphosate-resistant soybean and corn in 1996 and 1998, respectively, significantly reduced the application of PPO-inhibitors in crop fields [3]. Due to the widespread emergence of ALS-inhibitor and glyphosate resistance, and the slowly evolving nature of PPO-inhibitor resistance, PPO-inhibitors have recently increased in popularity [3-5]. Despite the long and widespread use of PPO-inhibitors, only eleven PPO-inhibitor-resistant weed species, including four *Amaranthus* species and two grass species, have been reported in six countries [3].

PPO-inhibiting herbicides hinder PPO enzyme function. There are two isoforms of the PPO enzyme-PPO1 (targeted to chloroplast) and PPO2 (mainly targeted to mitochondria, sometimes both chloroplasts and mitochondria), encoded by two nuclear genes *PPXI* and *PPXII*, respectively [3]. The PPO enzyme is crucial for the last step of heme and chlorophyll biosynthesis, namely the catalysis of protoporphyrinogen IX to protoporphyrin IX. PPO enzyme inhibition results in the accumulation of protoporphyrinogen IX in the chloroplasts or mitochondria, which leaks out to the cytosol where protoporphyrinogen IX gets oxidized to protoporphyrin IX. Protoporphyrin IX produces singlet reactive oxygen species in the presence of sunlight that disrupts the cell membrane and ultimately leads to plant death [5]. PPO-inhibitors include broadleaf, contact, and soil-applied herbicides. Resistance in weeds is conferred primarily by target site mutations in the *PPXII* gene [3, 6].

Different PPO-inhibitors chemistries are available, including heterocyclic phenyl ethers, oxadiazoles, phenyl imides, triazolinones, and pyrazoles [5]. The use of residual PPO-inhibitors, such as fomesafen (e.g., Flextar and Prefix), is increasing, particularly for weed control in soybean fields. Fomesafen is in the diphenyl ether class of PPO-inhibitors and can be applied pre-plant, pre-emergence, or post-emergence for the management of broadleaf weeds, grasses, and sedges in edible beans [5, 7]. Depending on conditions, the half-life of fomesafen ranges from six to twelve months in aerobic soil [7]. The application of residual PPO-inhibiting herbicide can impede the growth of the subsequent crop because of herbicide carryover injury.

The sensitivity of sorghum to herbicide residue in the soil from the previous crop (e.g. soybean) is of concern and constrains crop rotations. Sorghum was the most sensitive among common rotational crops such as corn, millet, and rice to fomesafen residues applied to beans [7]. Sorghum seedlings showed more than 40% phytotoxicity at 7 days after emergence in response to the PPO-inhibitor sulfentrazone [8]. The successful establishment of sorghum as a rotational crop with soybean requires sorghum cultivars to be tolerant to the herbicides applied to soybean. Thus, the development of herbicide-tolerant sorghum cultivars is critical for increasing sorghum production and expanding crop rotation options for growers. Recently, grain sorghum tolerant to ALS-inhibiting herbicide has been developed by introgressing the ALS-resistance gene from shatter cane, a weedy relative of sorghum, into sorghum through traditional breeding and is at the stage of commercialization [9]. However, there are not any commercial sorghum varieties tolerant to PPO-inhibiting herbicides. Identifying alleles conferring tolerance to PPO-inhibitors in sorghum will be useful for breeding PPO-inhibitor tolerant sorghum and hence, expanding the herbicide options for growers.

We observed fomesafen tolerance in a diverse sorghum population in the field and examined the underlying genetic mechanism of this tolerance. Our main goal was to examine fomesafen tolerance in sorghum. We performed a genome-wide association study (GWAS) in the sorghum biomass panel (SBP) to identify genomic regions associated with PPO inhibitor tolerance and examined the role of the *PPXI* gene in the observed tolerance using gene sequencing and expression analysis. We also developed a greenhouse assay to assess the sensitivity of sorghum lines to fomesafen and were able to reproduce the field phenotypes in the greenhouse. The result of our study will be useful for sorghum breeders to develop tolerant sorghum that avoids injury caused by residual PPO inhibitors.

## 2 Materials and Methods

### 2.1 Field Design and Phenotyping

The sorghum biomass panel (SBP) (n=718) was evaluated for residual herbicide injury during the 2015 field season at the Crop Sciences Research and Education Centers in Urbana, IL. The field was planted with soybeans in 2014 and sprayed with Flexstar, a member of the fomesafen class of PPO-inhibitors (group 14). The panel was planted in an augmented block design with a single replication. Check lines planted in each block included “Pacesetter”, “PRE0146”, “PRE0295”, “PRE0559”, “PRE0587”, “PRE1049”, “PRE1076”, and “SPX-901”. Each block consisted of 110 experimental treatments. Carryover injury was noticed approximately one month after planting and included blotches and chlorosis on the leaves. At this time, we assessed herbicide injury using a 1 to 9 scale, with 9 being the most sensitive [10].

### 2.2 Genotyping

A genotypic dataset (hereafter referred to as target set) scored using genotyping-by-sequencing was obtained using the procedures described by [11] and Thurber, Ma (12). In order to increase the marker density for the target panel, as described by Zhang, Fernandes (13). A genome-wide re-sequencing dataset (hereafter referred to as reference set) was used for imputing un-typed SNPs [14]. The reference panel was composed of 239 individuals and 5,512,653 SNPs anchored to the *Sorghum bicolor* reference genome version 3.1 (https://phytozome.jgi.doe.gov) [15]. The reference set data was filtered for heterozygosity (>10%), SNP coverage (<4X), and missing genotypes (>40%). Additionally, SNPs with minor allele count < 3 and depth < 3 were also filtered out before the imputation. The final reference panel used was composed of 239 individuals and 4,268,905 SNPs.

Before imputation, the target and reference panels were compared using conform-gt [16]. This step excluded target SNPs not present in the reference panel and adjusted the genomic position and chromosome strand to match the target and reference panels. Un-typed SNPs were imputed by chromosome, using option gt, window=80,000 bp, overlap=10,000 bp and ne=150,000. After filtering, Beagle version 4.1 was used to impute missing genotypes (option “gtgl”), followed by a phasing (option “gt”) step [17]. A window of 1500 bp and an overlap of 500 bp were used for both steps. The genotypic data were pruned based on linkage disequilibrium before conducting association analysis. The SNPs with an r^2^ value greater than 0.9 were removed with plink using a window size of 50 and a step size of 5 SNPs [18]. The markers were filtered for a minor allele frequency of 0.05 using GAPIT [19]. A total of 387,672 markers were included in the subsequent analyses.

### 2.3 Data analysis

The phenotypic data analysis was conducted in R (version 3.5.1) [20]. An analysis of variances (ANOVA) was performed to test the significant factors associated with the phenotypes observed in the field. The final linear mixed model, which included genotype, row, range and block, was run by using the “lme4” package in R [21]. Genotype was fit as a fixed effect, and block, range and row were fit as random effects. Best linear unbiased estimators (BLUEs) were calculated for each of the lines. The intercept was added to each line to get the final phenotypic data used for GWAS.

The genome-wide association study (GWAS) was conducted using the Genome Association and Prediction Integrated Tool (GAPIT) version 3.0 [19] in the R environment (version 3.5.1). The “CMLM” method was used to conduct the GWAS and a total of four principal components were included based on the scree plot. A false discovery rate of 0.05 was used to determine whether associations were significant [22].

### 2.4 Greenhouse assay

We selected ten representative sorghum lines based on their field phenotypes-five tolerant (PRE0278, PRE0282, PRE0520, PRE0545, PRE0546), and five sensitive (PRE0020, PRE0074, PRE0077, PRE0079, PRE0140) lines to develop the greenhouse assay. We initially determined the delimiting rate of pre-emergence fomesafen to differentiate herbicide tolerant and sensitive groups using the ten representative sorghum lines. Four replicates, with three plants per genotype per replicate, were planted in 1020 flats in the Plant Care Facility at the University of Illinois at Urbana-Champaign in Urbana in a randomized complete block design (RCBD) designed using the “agricolae” package in R (version 3.5.1) [23]. The seeds were pre-germinated in 100mm Petri dishes in the growth chamber before planting in the flats. Fomesafen was diluted at the log_3.16_ scale rates (0x, 0.001x, 0.003x, 0.01x, 0.03x, 0.1x and 0.316x) and uniformly sprayed in a spray chamber immediately after planting the stratified seeds in the pre-watered soil. The herbicide treated soil was covered with a layer of untreated soil to prevent any volatilization. Emergence counts were scored beginning at one day after treatment until five days after treatment. Herbicide injury severity was rated with a 1-9 scale described by Dear et al. (2003) weekly for three weeks after treatment. The fresh biomass was cut off and weighed at 21 days after the treatment. After weighing, the fresh biomass was dried for two days, and the dry weight (grams) was measured. The statistical analysis of the data was conducted in R (version 3.5.1) [20] using an ANOVA to determine the herbicide rate with the most significant difference between tolerant and susceptible lines. The ANOVA model included plot, replication, and sensitivity as factors. To confirm the delimiting rate identified from above assay, the greenhouse assay was repeated using ten sorghum lines (five sensitive and five tolerant) from the Sorghum Conversion Panel (SCP) [12] with five replications.

Using the rate determined during the preliminary experiments, we phenotyped a total of 100 sorghum lines from the SBP (50 sensitive and 50 tolerant). We included three replicates of each line in a randomized complete block design (RCBD) designed using the “agricolae” package in R with three plants per genotype in each replicate [23]. The lines were rated for the herbicide injury using the 1-9 scale described above. The statistical analysis of the data was conducted in R (version 3.5.1) [20] using an ANOVA test with the model that included plot, replication, and sensitivity.

### 2.5 PPXI as a candidate gene for herbicide tolerance in sorghum

#### 2.5.1 Sequence variant detection

The ten representative sorghum lines, along with an additional six sorghum lines (four sensitive and two tolerant) from the SBP panel, were surveyed for sequence variation in exonic regions of *PPXI*. The phenotype of these 16 lines was consistent between field and greenhouse studies. Fresh leaf tissue was collected from three-week-old sorghum plants and immediately placed in liquid nitrogen. Four primers (Table S1) were designed to amplify the cDNA sequences in the chloroplastic *PPXI* gene region (Accession no. XM_002455439.2) using NCBI Primer-BLAST software (https://www.ncbi.nlm.nih.gov/tools/primer-blast/). RNA was extracted using TriZol (ThermoFisher Scientific, Waltham, MA) and cleaned with a Qiagen RNAeasy miniElute cleanup kit (QIAGEN, Germantown, MD) as described in Fall, Salazar (24). The cDNA was synthesized from mRNA using revert aid first strand cDNA synthesis kit (ThermoFisher Scientific, Waltham, MA) and a random hexamer using the manufacturer’s protocol.

The cDNA amplification was performed in a 25 µl reaction containing 0.6 U of dreamTaq polymerase (ThermoFisher Scientific, Waltham, MA), 1× dreamTaq green buffer, 0.2 mM dNTP, 0.4 μm of each the forward and reverse primers, 0.5mM MgCl_2_, and nuclease-free water in a thermocycler using three-step cycling. One initial cycle of denaturation at 95°C for 2 min was carried out; followed by 35 cycles of denaturation at 95°C for 30 s; annealing at 52°C for the 30s; and extension at 72°C for 1 min; and a final cycle of extension at 72°C for 10 min. The amplification of the GC-rich region (primer 1) was performed with high fidelity Q5 polymerase (New England Biolabs Inc, Ipswich, MA) using the manufacturer’s protocol. The PCR products were confirmed by running agarose gel electrophoresis on a 1.0 % agarose gel. The gel image was visualized using a UVP GelDoc-It2 310 imager (UVP, Upland, CA). The positive PCR products were cleaned using Wizard® SV Gel and PCR Clean-Up System (Promega Corporation, Madison, WI) and submitted for Sanger sequencing at Roy J. Carver Biotechnology Center at the University of Illinois at Urbana-Champaign.

The sequences were trimmed and aligned with the *PPXI* mRNA sequence from the sorghum reference (Accession no. XM_002455439) using MUSCLE within Molecular Evolutionary Genetics Analysis (MEGA) software version 7.0 and default parameters [25]. Individual insert sequences for each primer pair set were concatenated to obtain a full sequence of *PPXI*, and overlapping regions were merged. The gene sequences were compared among tolerant and susceptible lines, along with the reference BTx623 *PPXI* allele, to determine the variation present within the *PPXI* gene.

#### 2.5.2 Gene expression analysis

The ten representative sorghum lines were assayed for *PPXI* expression using quantitative reverse transcription PCR (qRT-PCR). Seed for the lines were planted in 1020 flats with three replicates arranged in a randomized complete block design (RCBD). Fomasafen was applied using the method described above. Leaf samples were collected into liquid nitrogen seventeen days after herbicide treatment. The herbicide injury level was rated using the 1-9 scale described by Dear et al. (2003) before sample collection.

We extracted the RNA as described above. To test RNA integrity and for DNA contamination, the RNA was run out on a 1% gel using electrophoresis. The primers and probes for TaqMan® gene expression assay were designed using Integrated DNA Technology (IDT) PrimerQuest Tool (https://www.idtdna.com/PrimerQuest/Home/Index) according to the IDT guidelines and synthesized by ThermoFisher. The specificity of primers and probes were checked using Primer Blast. The combinations of primers and probes that resulted in a product specific to *PPXI* gene were retained (Table S2).

One-step qRT-PCR was performed in an ABI Prism 7000 detection system (Applied Biosystems) with equal RNA concentrations. A total reaction volume of 20µl using Verso 1-step RT-qPCR ROX Mix kit (ThermoFisher Scientific, Waltham, MA) according to the manufacture’s protocol was used. The final concentration of primers and probe in the reaction were 450nM and 125nM, respectively. The amplification program consisted of one cycle of cDNA synthesis at 50°C for 15 min; one cycle of thermo-start polymerase activation at 95°C for 15 min; 40 cycles of denaturation at 95°C for 15s and annealing/extension at 60°C for 60s. The *PP2A* gene was used as an internal reference gene for data normalization, as suggested by Sudhakar Reddy, Srinivas Reddy (26). The efficiency of both *PPXI* and *PP2A* Taqman assays were tested using a qPCR standard curve using a 10-fold serial dilution of RNA with final concentration ranging from 2pg/µl to 20,000pg/µl. The formula used to calculate assay efficiency was as follows:

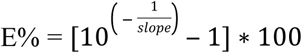

Ct values were determined based on three technical replicates of each sample, and mean Ct values for the sensitive and tolerant groups were calculated. The mean Ct values for both groups were transformed into relative quantification (RQ) using the Pffalfl method [27].

## 3 Results

### 3.1 Evaluation of herbicide injury in the field

The SBP panel showed a wide range of herbicide injury ratings, ranging from 1 to 9, with a mean rating of 4.2 and a standard deviation of 1.9 (Fig. 1). The tolerant lines were without any symptoms, while sensitive lines showed leaf blotches and chlorosis. The phenotypes were continuously distributed. The line effects were highly significant and explained the largest proportion of phenotypic variation among range, row, and block (Table 1).

**Table 1.** Analysis of variance (ANOVA) of herbicide injury data of sorghum biomass panel (SBP) obtained from the 2015 field season and SBP subset (100 sorghum lines) from the greenhouse assay.

| Field Assay |  |  |  | Greenhouse Assay |  |  |  |  |  |
| --- | --- | --- | --- | --- | --- | --- | --- | --- | --- |
|  |  |  |  | Week 2 |  |  | Week 3 |  |  |
| Source | MS | F | Pr > F | MS | F | Pr > F | MS | F | Pr > F |
| Lines | 3.8054 | 7.4801 | 2.00E-16 *** a | 4.9 | 1.45 | 0.0237* | 5.6 | 1.54 | 0.0103* |
| Rep | NA<sup>b</sup> | NA | NA | 1.8 | 0.54 | 0.582 | 22 | 6.01 | 0.0032** |
| Range | 0.8359 | 1.6431 | 0.033* | NA | NA | NA | NA | NA | NA |
| Row | 0.8863 | 1.7421 | 0.04237* | NA | NA | NA | NA | NA | NA |
| Block | 0.5485 | 1.0783 | 0.3858 | NA | NA | NA | NA | NA | NA |
| Residual | 0.5087 |  |  | 3.4 |  |  | 3.7 |  |  |

<sup>a</sup> ‘\*\*\*’ denotes p - value < 0.001, ‘\*\*’ denotes p - value < 0.01, ‘\*’ denotes p - value < 0.05
<sup>b</sup> NA indicates not available.

**Fig. 1.** Phenotypic distribution (A) and phenotypes (B and C) of herbicide injury in the sorghum biomass panel during the 2015 field season.

### 3.2 Phenotype confirmation in the greenhouse

Herbicide injury, assessed using visual ratings, was significantly different between tolerant and sensitive groups at the 0.1x rate for every rating (each week for three weeks post spray) (Fig. S1). The delimiting rate of 0.1x was further confirmed on five sensitive and five tolerant lines from the SCP in a subsequent greenhouse assay. Significant differences were observed between the sensitive and tolerant groups at the second and third weeks after the herbicide treatment in the SCP (Fig. S1). For the seven herbicide application rates that were tested, no significant differences between sensitive and tolerant groups for any rate were observed for emergence counts, dry weight, and fresh weight. Thus, visual ratings of herbicide injury were used for further assessment of herbicide tolerance.

We evaluated 100 sorghum lines from the SBP (50 sensitive and 50 tolerant) for pre-emergence herbicide injury in the greenhouse to validate the field findings. In the resulting ANOVA model, genotype was significant two and three weeks after the herbicide treatment (Table 1). Significant differences were also observed between sensitive and tolerant groups, confirming the field phenotypes (Fig. S1).

### 3.3 SNPs significantly associated with herbicide tolerance

A total of 26 SNPs were significant in the GWAS at an FDR of 5% (Table 2). All the significant SNPs were located on chromosome 3 in the region from 12,937,584 bp to 13,152,838 bp (Fig. 2). This 215.3 kb region encompasses eight genes and significant linkage disequilibrium (LD) was found in this region (Fig. 3). Among 26 significant SNPs, six were genic and within four unique genes-Sobic.003G136200 (germin-like protein), Sobic.003G136500 (not annotated), Sobic.003G136800 (SNF7 family protein), and Sobic.003G136900 (phytochrome interacting factor 3). The ten most significant SNPs included eight SNPs in intergenic regions that were close to Sobic.003G136800 (SNF7 family protein), Sobic.003G137000 (RING/U-box superfamily protein), Sobic.003G136900 (phytochrome interacting factor 3) and Sobic.003G137100 (*PPXI*). The significant SNP S03_13152838 (p< 0.001) was located 12kb upstream of the *PPXI* gene.

**Table 2.**
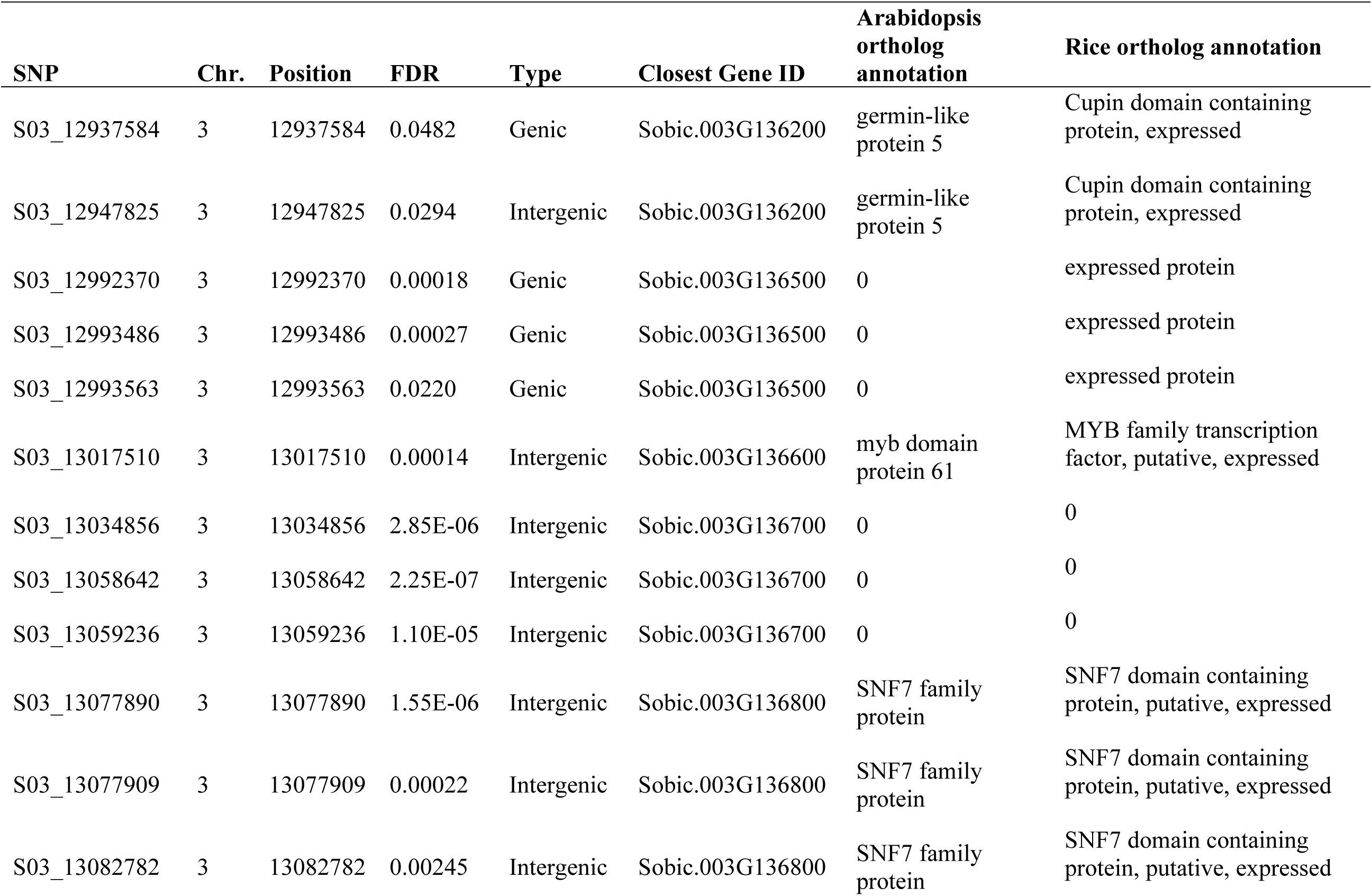
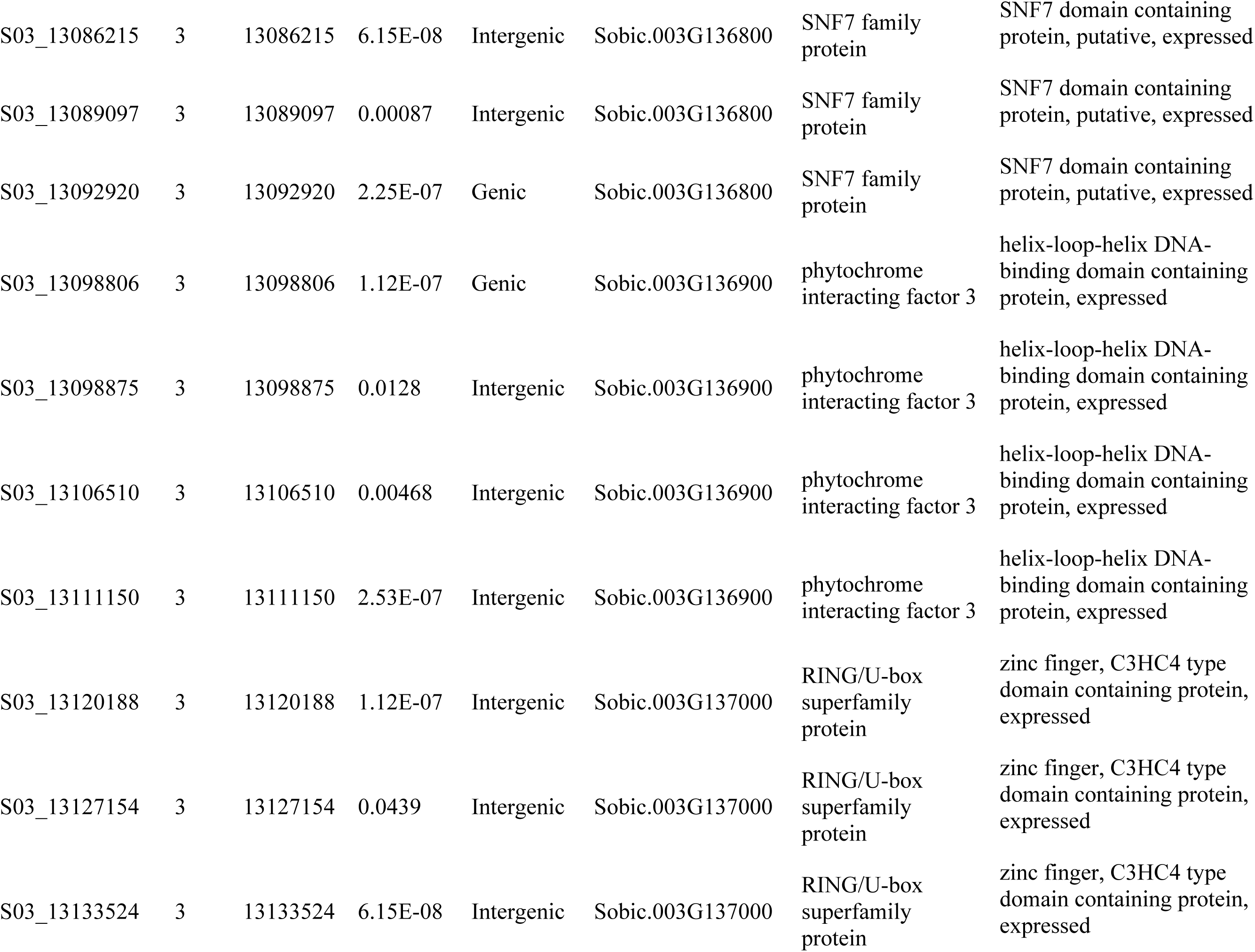
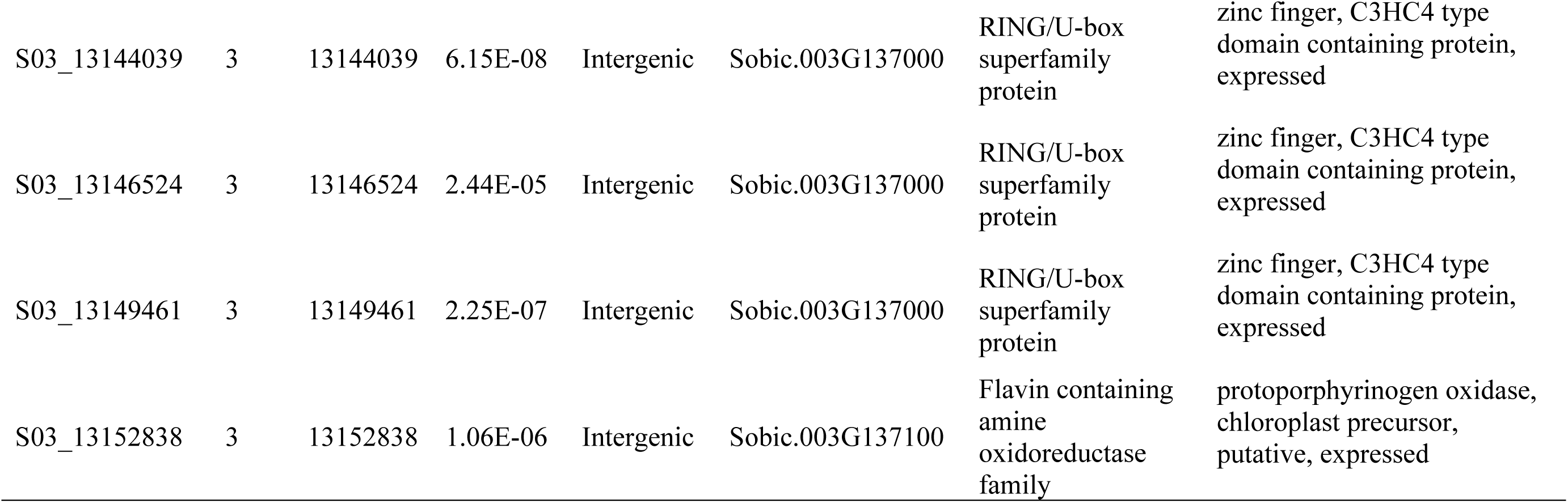
Significant SNPs associated with the herbicide tolerance based on the genome-wide association study.

| SNP | Chr. | Position | FDR | Type | Closest Gene ID | Arabidopsis ortholog annotation | Rice ortholog annotation |
| --- | --- | --- | --- | --- | --- | --- | --- |
| S03_12937584 | 3 | 12937584 | 0.0482 | Genic | Sobic.003G136200 | germin-like protein 5 | Cupin domain containing protein, expressed |
| S03_12947825 | 3 | 12947825 | 0.0294 | Intergenic | Sobic.003G136200 | germin-like protein 5 | Cupin domain containing protein, expressed |
| S03_12992370 | 3 | 12992370 | 0.00018 | Genic | Sobic.003G136500 | 0 | expressed protein |
| S03_12993486 | 3 | 12993486 | 0.00027 | Genic | Sobic.003G136500 | 0 | expressed protein |
| S03_12993563 | 3 | 12993563 | 0.0220 | Genic | Sobic.003G136500 | 0 | expressed protein |
| S03_13017510 | 3 | 13017510 | 0.00014 | Intergenic | Sobic.003G136600 | myb domain protein 61 | MYB family transcription factor, putative, expressed |
| S03_13034856 | 3 | 13034856 | 2.85E-06 | Intergenic | Sobic.003G136700 | 0 | 0 |
| S03_13058642 | 3 | 13058642 | 2.25E-07 | Intergenic | Sobic.003G136700 | 0 | 0 |
| S03_13059236 | 3 | 13059236 | 1.10E-05 | Intergenic | Sobic.003G136700 | 0 | 0 |
| S03_13077890 | 3 | 13077890 | 1.55E-06 | Intergenic | Sobic.003G136800 | SNF7 family protein | SNF7 domain containing protein, putative, expressed |
| S03_13077909 | 3 | 13077909 | 0.00022 | Intergenic | Sobic.003G136800 | SNF7 family protein | SNF7 domain containing protein, putative, expressed |
| S03_13082782 | 3 | 13082782 | 0.00245 | Intergenic | Sobic.003G136800 | SNF7 family protein | SNF7 domain containing protein, putative, expressed |
| S03_13086215 | 3 | 13086215 | 6.15E-08 | Intergenic | Sobic.003G136800 | SNF7 family protein | SNF7 domain containing protein, putative, expressed |
| S03_13089097 | 3 | 13089097 | 0.00087 | Intergenic | Sobic.003G136800 | SNF7 family protein | SNF7 domain containing protein, putative, expressed |
| S03_13092920 | 3 | 13092920 | 2.25E-07 | Genic | Sobic.003G136800 | SNF7 family protein | SNF7 domain containing protein, putative, expressed |
| S03_13098806 | 3 | 13098806 | 1.12E-07 | Genic | Sobic.003G136900 | phytochrome interacting factor 3 | helix-loop-helix DNA-binding domain containing protein, expressed |
| S03_13098875 | 3 | 13098875 | 0.0128 | Intergenic | Sobic.003G136900 | phytochrome interacting factor 3 | helix-loop-helix DNA-binding domain containing protein, expressed |
| S03_13106510 | 3 | 13106510 | 0.00468 | Intergenic | Sobic.003G136900 | phytochrome interacting factor 3 | helix-loop-helix DNA-binding domain containing protein, expressed |
| S03_13111150 | 3 | 13111150 | 2.53E-07 | Intergenic | Sobic.003G136900 | phytochrome interacting factor 3 | helix-loop-helix DNA-binding domain containing protein, expressed |
| S03_13120188 | 3 | 13120188 | 1.12E-07 | Intergenic | Sobic.003G137000 | RING/U-box superfamily protein | zinc finger, C3HC4 type domain containing protein, expressed |
| S03_13127154 | 3 | 13127154 | 0.0439 | Intergenic | Sobic.003G137000 | RING/U-box superfamily protein | zinc finger, C3HC4 type domain containing protein, expressed |
| S03_13133524 | 3 | 13133524 | 6.15E-08 | Intergenic | Sobic.003G137000 | RING/U-box superfamily protein | zinc finger, C3HC4 type domain containing protein, expressed |
| S03_13144039 | 3 | 13144039 | 6.15E-08 | Intergenic | Sobic.003G137000 | RING/U-box superfamily protein | zinc finger, C3HC4 type domain containing protein, expressed |
| S03_13146524 | 3 | 13146524 | 2.44E-05 | Intergenic | Sobic.003G137000 | RING/U-box superfamily protein | zinc finger, C3HC4 type domain containing protein, expressed |
| S03_13149461 | 3 | 13149461 | 2.25E-07 | Intergenic | Sobic.003G137000 | RING/U-box superfamily protein | zinc finger, C3HC4 type domain containing protein, expressed |
| S03_13152838 | 3 | 13152838 | 1.06E-06 | Intergenic | Sobic.003G137100 | Flavin containing amine oxidoreductase family | protoporphyrinogen oxidase, chloroplast precursor, putative, expressed |

**Fig. 2.** Manhattan plot for the genome-wide association mapping. Twenty-six significant SNPs were detected on chromosome 3.

**Fig. 3.** Linkage disequilibrium (LD) plot for the significant SNPs in the 215 kb region of chromosome 3. The Manhattan plot for the region is shown above and the linkage disequilibrium shown below. The Manhattan plot included only the significant SNPs from the association analysis. In the LD plot, the r^2^ values between significant SNPs are shown. Red indicates high amounts of linkage disequilibrium, while yellow indicates low linkage disequilibrium.

We hypothesized that *PPXI* may not have had any significant genic associations because there were no SNPs within the gene in the association mapping genotypic dataset. Thus, we examined whether there were variants in the *PPXI* gene included in the analysis. There were five SNPs in the *PPXI* gene in genotypic data, three of them (S03_13165379, S03_13170697, and S03_13170922) were in exons, and two of them (S03_13165710 and S03_13169856) were in introns. None of these SNPs were significant in the association analysis.

### 3.4 Examining *PPXI* gene as a possible candidate

Because PPO enzymes are targeted by the herbicide, and due to the gene’s proximity to significant SNPs in the GWAS, we examined *PPXI* as a candidate gene. Because our dataset was not exhaustive in terms of sequence variants in the panel, we conjectured that we might be missing important functional variation in the *PPXI* gene in our genotypic dataset. We hypothesized that variation in the active site of the enzyme might underlie the herbicide tolerance we observed. A total of 16 lines (seven tolerant and nine sensitive) from the SBP were selected to survey the variation in the *PPXI* gene sequences. We obtained sequencing data for 12 of those lines, while the remaining four lines (both sensitive and tolerant) could not be examined because of poor sequencing quality. We did not detect any sequence variation in the *PPXI* mRNA region. We concluded that *PPXI* is highly conserved in sorghum, and sequence variation in the *PPXI* gene does not underlie the herbicide tolerance we observed.

In light of the lack of sequence variation, we hypothesized that the *PPXI* gene might be differentially expressed between tolerant and sensitive lines. Lermontova and Grimm (28) reported that overexpression of the wild-type Arabidopsis *PPXI* gene in transgenic tobacco resulted in a five-fold increase in enzymatic activity, which prevented the accumulation of toxic protoporphyrinogen IX and, thus, increased tolerance to the PPO-inhibitor acifluorfen. Therefore, we selected a total of 10 lines to examine *PPXI* expression after fomesafen application. Tolerant and sensitive groups selected for the gene expression study showed significant phenotypic differences in the greenhouse when the tissue was collected (*p*<0.0001). However, there were no significant differences in *PPXI* gene expression between tolerant and sensitive groups (RQ = 1.27). It is unlikely that *PPXI* underlies the herbicide tolerance we observed.

## 4 Discussion

Herbicide resistance mechanisms can be classified into two categories: target site resistance (TSR) and non-target site resistance (NTSR). TSR includes genic mutations that result in structural changes in the proteins targeted by the herbicide, which then reduces herbicide binding [29]. Alternately, NTSR includes diverse mechanisms, including reduced herbicide uptake/translocation, increased herbicide detoxification, decreased herbicide activation rates, and herbicide sequestration [30]. Metabolism-based NTSR is associated with the herbicide detoxification due to the increased activity of enzyme complexes, including esterases, cytochrome P450s, glutathione S-transferase (GSTs), and UDP-glucosyl transferase. Unlike TSR, metabolism-based NTSR is largely polygenic and confers resistance to herbicides with multiple modes of action [30].

*PPO1* and *PPO2* are both molecular targets of PPO-inhibiting herbicides. In weeds, several mutations in *PPXII*, which lead to TSR, have been reported. For example, a mutation involving the loss of three nucleotides in the coding sequence of *PPXII* conferred resistance to a PPO-inhibiting herbicide in waterhemp (*Amaranthus tuberculatus*) [31]. This mutation resulted in the deletion of a glycine residue at the 210^th^ position (ΔG210)) of the PPO2 enzyme [31, 32]. In Palmer amaranth (*Amaranthus palmeri*), the substitution of arginine to glycine/methionine at the 128^th^ position of the PPO2 enzyme was observed in addition to the ΔG210 mutation in fomesafen-resistant weeds [33, 34]. Recently, another mutation, e.g., the substitution of glycine to alanine in the catalytic domain of *PPXII* at position 399 (G_399_), was reported in Palmer amaranth resistant to PPO-inhibitors [35]. In common ragweed (*Ambrosia artemisiifolia)*, a mutation causing the substitution of an arginine (Arg98) for a leucine codon at the R98L position of the PPO2 enzyme conferred resistance to a PPO-inhibitor [36]. It is important to note that all of these mutations in weeds that conferred herbicide resistance occurred in *PPXII*. However, Lermontova and Grimm (28) showed that overexpression of *PPXI* from wild-type Arabidopsis increased the tolerance to the PPO-inhibitor acifluorfen in tobacco.

In the GWAS, we identified a significant SNP on chromosome 3 that was 12kb upstream of *PPXI*. We hypothesized that *PPXI* might be responsible for the observed herbicide tolerance in the SBP. Furthermore, we hypothesized that the lack of detection of a SNP in the genic region was due to the incomplete nature of the genotypic dataset in the *PPXI* region. However, we could find neither sequence variation in the coding region of *PPXI*, nor an expression difference between tolerant and sensitive lines. In the genotypic data used for GWAS, there were only three SNPs (S03_13165379, S03_13170697, and S03_13170922) and five haplotypes in the *PPXI* exonic regions. A total of 694 out of 718 lines from the SBP had the same haplotype as the reference line. For the SNPs S03_13165379 and S03_13170697, thirteen common lines out of 718 lines had alternate alleles, and for the SNP S03_13170922, eleven out of 718 lines had alternate alleles. This suggests that *PPXI* is conserved and that the herbicide tolerance observed in the sorghum population might be related to NTSR, especially metabolism-based resistance. In addition, if it were a target site mutation with little environmental influence, we would have expected a bimodal distribution with our field phenotypes. However, the observed distribution indicates that there were strong environmental effects or that multiple genes are responsible for the herbicide tolerance observed in our population. Our hypothesis of multiple genes underlying the observed phenotype is also supported by the LD structure at the regions that were significantly associated with herbicide tolerance.

Non-target site resistance has been observed in several crop and weed species. In soybean, the natural tolerance to diphenyl ether class of PPO-inhibitors is due to the rapid metabolic cleavage of diphenyl ether bond [37] and homoglutathione conjugation is involved in the detoxification of diphenyl ether [38]. Similarly, rapid glutathione conjugation also conferred tolerance to diphenyl ether class of PPO-inhibitors in peas [39]. In soybean, metabolism-based tolerance was observed to the pre-application of sulfentrazone herbicide, another class of PPO-inhibiting herbicide [40]. The degradation of sulfentrazone was due to the oxidation of the methyl group on the triazolinone ring [40]. In Palmar amaranth, some plants resistant to fomesafen did not have a target site mutation in *PPXII*, which suggests that these plants might be presenting NTSR [5].

We investigated other plausible genes from the GWAS for a potential role in metabolism based NTSR. The most significant SNPs from the GWAS were in the intergenic region proximal to Sobic.003G137000 (RING/U-box superfamily protein) and Sobic.003G136800 (SNF7 family protein). We found significant SNPs in the genic regions of Sobic.003G136200 (germin-like protein), Sobic.003G136800 (SNF7 family protein), and Sobic.003G136900 (phytochrome interacting factor 3).

Based on the literature, the RING/U-box superfamily protein-encoding gene is a strong candidate. Mahmood et al. (2016) identified a cis-regulatory motif involved in the formation of a CUL4-RING ubiquitin ligase complex and zinc finger transcription factors regulating herbicide metabolism related (HMR) genes such as cytochrome P450s, nitronate monoxygenase, and glutathione S-transferase in *Arabidopsis* and rice. The zinc finger transcription factors had a similar level of expression as that of HMR genes and were highly expressed in response to herbicides [41]. The zinc finger protein has also been reported to negatively regulate plant cell death in *Arabidopsis* [42]. In our study, three significant SNPs were detected downstream and three upstream of the gene Sobic.003G137000 (RING/U-box superfamily protein). The absence of a significant SNP in the gene region might be due to the absence of SNPs in this gene region in the genotypic dataset. It is plausible that the herbicide resistance observed in our population might be associated with the zinc finger domain regulated HMR resistance.

There were five SNPs in the intergenic region and one SNP in the genic region of Sobic.003G136800, which encodes a SNF-7 family protein. SNF-7 proteins are part of endosomal sorting complexes required for transport (ESCRT) machinery that is involved in multivesicular body biogenesis and sorting of ubiquitinated membrane proteins for degradation [43]. The SNF-7 gene could be involved in the vacuolar sorting of proteins targeted by metabolism-related genes for degradation.

In conclusion, we identified PPO-inhibitor tolerance in a diverse sorghum population. We developed a greenhouse assay to test for fomesafen tolerance in sorghum and confirmed field phenotypes. We identified a region of chromosome 3 that encompassed nine genes as being associated with fomesafen tolerance. We found that *PPXI* is highly conserved in sorghum and likely does not underlie the observed herbicide tolerance. Instead, the mechanism underlying this tolerance might be metabolism-based resistance, possibly regulated by the action of multiple genes, as indicated by continuous phenotypic distribution and LD structure within the region. Further experiments will confirm the role of candidate genes. The overall results of our study will be useful for sorghum breeders to develop fomasafen tolerant sorghum that avoids injury caused by residual PPO inhibitors and enable more diversified crop rotations.

## 6 Supporting information captions

**Fig. S1.** Greenhouse assay for phenotypic evaluation. Phenotypic differences between 10 sensitive and tolerant representative sorghum lines selected from the sorghum biomass panel (SBP) for seven herbicide rates at the first week (A), second week (B), and third week (C) after herbicide treatment. Sensitive and tolerant groups were significant at the rate 0.1x. Significant phenotypic differences (*p<0*.*0001*) were observed in the subset of 10 sorghum lines from sorghum conversion panel (SCP) (D), and 100 sorghum lines from the SBP (E) with the 0.1x rate of herbicide at second and third week after the treatment.

## References

1. Besançon T, Heiniger R, Weisz R, Everman W. Weed response to agronomic practices and herbicide strategies in grain sorghum. Agronomy Journal. 2017;109(4):1642–50.

2. Barber T, Scott B, Norsworthy JK. Weed control in grain sorghum. Arkansas Grain Sorghum Production Handbook. 2015;Chapter 8:1-14.

3. Dayan FE, Barker A, Tranel PJ. Origins and structure of chloroplastic and mitochondrial plant protoporphyrinogen oxidases: implications for the evolution of herbicide resistance. Pest management science. 2018;74(10):2226–34.

4. Cornelius CD, Bradley KW. Carryover of Common Corn and Soybean Herbicides to Various Cover Crop Species. Weed Technology. 2017;31(1):21–31. doi:https://doi.org/10.1614/WT-D-16-00062.1.

5. Salas RA, Burgos NR, Tranel PJ, Singh S, Glasgow L, Scott RC, et al. Resistance to PPO-inhibiting herbicide in Palmer amaranth from Arkansas. Pest management science. 2016;72(5):864–9.

6. Thinglum KA, Riggins CW, Davis AS, Bradley KW, Al-Khatib K, Tranel PJ. Wide distribution of the waterhemp (Amaranthus tuberculatus) ΔG210 PPX2 mutation, which confers resistance to PPO-inhibiting herbicides. Weed science. 2011;59(1):22–7.

7. Cobucci T, Prates HT, Falcão CL, Rezende MM. Effect of imazamox, fomesafen, and acifluorfen soil residue on rotational crops. Weed Science. 1998;46(2):258–63.

8. Concenço G, Andres A, Schreiber F, Palharini WG, Martins MB, Moisinho IS, et al. Sweet Sorghum Establishment after Application of Residual Herbicides. Embrapa Clima Temperado-Artigo em periódico indexado (ALICE). 2018.

9. Werle R, Tenhumberg B, Lindquist JL. Modeling shattercane dynamics in herbicide-tolerant grain sorghum cropping systems. Ecological modelling. 2017;343:131–41.

10. Dear B, Sandral G, Spencer D, Khan M, Higgins T. The tolerance of three transgenic subterranean clover (Trifolium subterraneum L.) lines with the bxn gene to herbicides containing bromoxynil. Australian Journal of Agricultural Research. 2003;54(2):203–10.

11. Fernandes SB, Dias KO, Ferreira DF, Brown PJ. Efficiency of multi-trait, indirect, and trait-assisted genomic selection for improvement of biomass sorghum. Theoretical and applied genetics. 2018;131(3):747–55.

12. Thurber CS, Ma JM, Higgins RH, Brown PJ. Retrospective genomic analysis of sorghum adaptation to temperate-zone grain production. Genome Biol. 2013;14(6):R68. Epub 2013/06/28. doi:10.1186/gb-2013-14-6-r68. PubMed PMID: 23803286; PubMed Central PMCID: PMCPMC3706989.

13. Zhang X, Fernandes SB, Kaiser C, Adhikari P, Brown PJ, Mideros SX, et al. Conserved defense responses between maize and sorghum to Exserohilum turcicum. BMC Plant Biol. 2020;20(1):67. Epub 2020/02/12. doi:10.1186/s12870-020-2275-z. PubMed PMID: 32041528; PubMed Central PMCID: PMCPMC7011368.

14. Valluru R, Gazave EE, Fernandes SB, Ferguson JN, Lozano R, Hirannaiah P, et al. Deleterious mutation burden and its association with complex traits in sorghum (*Sorghum bicolor*). Genetics. 2019;211(3):1075–87. Epub 2019/01/10. doi:10.1534/genetics.118.301742. PubMed PMID: 30622134; PubMed Central PMCID: PMCPMC6404259.

15. McCormick RF, Truong SK, Sreedasyam A, Jenkins J, Shu S, Sims D, et al. The *Sorghum bicolor* reference genome: improved assembly, gene annotations, a transcriptome atlas, and signatures of genome organization. Plant J. 2018;93(2):338–54. Epub 2017/11/22. doi:10.1111/tpj.13781. PubMed PMID: 29161754.

16. Browning SR, Browning BL. Rapid and accurate haplotype phasing and missing-data inference for whole-genome association studies by use of localized haplotype clustering. Am J Hum Genet. 2007;81(5):1084–97. Epub 2007/10/10. doi:10.1086/521987. PubMed PMID: 17924348; PubMed Central PMCID: PMCPMC2265661.

17. Browning BL, Browning SR. Genotype imputation with millions of reference samples. Am J Hum Genet. 2016;98(1):116–26. Epub 2016/01/11. doi:10.1016/j.ajhg.2015.11.020. PubMed PMID: 26748515; PubMed Central PMCID: PMCPMC4716681.

18. Purcell S, Neale B, Todd-Brown K, Thomas L, Ferreira MA, Bender D, et al. PLINK: a tool set for whole-genome association and population-based linkage analyses. The American journal of human genetics. 2007;81(3):559–75.

19. Tang Y, Liu X, Wang J, Li M, Wang Q, Tian F, et al. GAPIT Version 2: An Enhanced Integrated Tool for Genomic Association and Prediction. Plant Genome-Us. 2016;9(2). Epub 2016/11/30. doi:10.3835/plantgenome2015.11.0120. PubMed PMID: 27898829.

20. R Core Team. A language and environment for statistical computing. Vienna, Austria: R Foundation for Statistical Computing; 2012. URL https://www.R-projectorg. 2019.

21. Bates D, Sarkar D, Bates MD, Matrix L. The lme4 package. R package version. 2007;2(1):74.

22. Benjamini Y, Hochberg Y. Controlling the false discovery rate: a practical and powerful approach to multiple testing. Journal of the Royal statistical society: series B (Methodological). 1995;57(1):289–300.

23. de Mendiburu F, de Mendiburu MF. Package ‘agricolae’. R Package, Version. 2019:1.2–1.

24. Fall LA, Salazar MM, Drnevich J, Holmes JR, Tseng M-C, Kolb FL, et al. Field pathogenomics of Fusarium head blight reveals pathogen transcriptome differences due to host resistance. Mycologia. 2019:1–11.

25. Kumar S, Stecher G, Tamura K. MEGA7: molecular evolutionary genetics analysis version 7.0 for bigger datasets. Molecular biology and evolution. 2016;33(7):1870–4.

26. Sudhakar Reddy P, Srinivas Reddy D, Sivasakthi K, Bhatnagar-Mathur P, Vadez V, Sharma KK. Evaluation of sorghum [Sorghum bicolor (L.)] reference genes in various tissues and under abiotic stress conditions for quantitative real-time PCR data normalization. Frontiers in plant science. 2016;7:529.

27. Pfaffl MW. A new mathematical model for relative quantification in real-time RT–PCR. Nucleic acids research. 2001;29(9):e45–e.

28. Lermontova I, Grimm B. Overexpression of plastidic protoporphyrinogen IX oxidase leads to resistance to the diphenylether herbicide acifluorfen. Plant Physiology. 2000;122(1):75–84.

29. Délye C, Duhoux A, Pernin F, Riggins CW, Tranel PJ. Molecular mechanisms of herbicide resistance. Weed Science. 2015;63(SP1):91–115.

30. Chahal PS, Aulakh JS, Jugulam M, Jhala AJ. Herbicide-resistant Palmer amaranth (Amaranthus palmeri S. Wats.) in the United States—mechanisms of resistance, impact, and management. Herbicides, Agronomic Crops and Weed Biology Rijeka, Croatia: InTech. 2015:1–29.

31. Patzoldt WL, Hager AG, McCormick JS, Tranel PJ. A codon deletion confers resistance to herbicides inhibiting protoporphyrinogen oxidase. Proceedings of the National Academy of Sciences. 2006;103(33):12329–34.

32. Lee RM, Hager AG, Tranel PJ. Prevalence of a novel resistance mechanism to PPO-inhibiting herbicides in waterhemp (Amaranthus tuberculatus). Weed science. 2008;56(3):371–5.

33. Salas-Perez RA, Burgos NR, Rangani G, Singh S, Refatti JP, Piveta L, et al. Frequency of Gly-210 deletion mutation among protoporphyrinogen oxidase inhibitor–resistant Palmer amaranth (Amaranthus palmeri) populations. Weed science. 2017;65(6):718–31.

34. Varanasi VK, Brabham C, Norsworthy JK, Nie H, Young BG, Houston M, et al. A statewide survey of PPO-inhibitor resistance and the prevalent target-site mechanisms in Palmer amaranth (Amaranthus palmeri) accessions from Arkansas. Weed science. 2018;66(2):149–58.

35. Rangani G, Salas-Perez RA, Aponte RA, Knapp M, Craig IR, Meitzner T, et al. A Novel Single-Site Mutation in the Catalytic Domain of Protoporphyrinogen Oxidase IX (PPO) Confers Resistance to PPO-Inhibiting Herbicides. Frontiers in plant science. 2019;10:568.

36. Rousonelos SL, Lee RM, Moreira MS, VanGessel MJ, Tranel PJ. Characterization of a common ragweed (Ambrosia artemisiifolia) population resistant to ALS-and PPO-inhibiting herbicides. Weed science. 2012;60(3):335–44.

37. Frear D, Swanson H, Mansager E. Acifluorfen metabolism in soybean: diphenylether bond cleavage and the formation of homoglutathione, cysteine, and glucose conjugates. Pesticide Biochemistry and Physiology. 1983;20(3):299–310.

38. Skipsey M, Andrews CJ, Townson JK, Jepson I, Edwards R. Substrate and thiol specificity of a stress-inducible glutathione transferase from soybean. FEBS letters. 1997;409(3):370–4.

39. Edwards R. Characterisation of glutathione transferases and glutathione peroxidases in pea (Pisum sativum). Physiologia Plantarum. 1996;98(3):594–604.

40. Dayan FE, Weete JD, Duke SO, Hancock HG. Soybean (Glycine max) cultivar differences in response to sulfentrazone. Weed Science. 1997;45(5):634–41.

41. Mahmood K, Mathiassen SK, Kristensen M, Kudsk P. Multiple herbicide resistance in Lolium multiflorum and identification of conserved regulatory elements of herbicide resistance genes. Frontiers in Plant Science. 2016;7:1160.

42. Dietrich RA, Richberg MH, Schmidt R, Dean C, Dangl JL. A novel zinc finger protein is encoded by the Arabidopsis LSD1 gene and functions as a negative regulator of plant cell death. Cell. 1997;88(5):685–94.

43. Cui Y, He Y, Cao W, Gao J, Jiang L. The multivesicular body and autophagosome pathways in plants. Frontiers in plant science. 2018;9.

